# Similarities and differences in molecular epidemiology of third-generation cephalosporin-resistant *Escherichia coli* carried by dogs living in urban and nearby rural settings and associated behavioural risk factors

**DOI:** 10.1101/2022.04.23.489260

**Authors:** Jordan E. Sealey, Ashley Hammond, Oliver Mounsey, Virginia C. Gould, Kristen K. Reyher, Matthew B. Avison

## Abstract

**Objectives:** Our aims were to compare faecal third-generation cephalosporin-resistant (3GC-R) *Escherichia coli* isolates from dogs living in a city and in a rural area ~30 km away; to compare isolates from dogs, cattle, and humans in these regions; to determine risk factors associated with 3GC-R *E. coli* carriage in these two cohorts of dogs.

**Methods:** 600 dogs were included, with faecal samples processed to recover 3GC-R *E. coli* using 2 mg/L cefotaxime. WGS was by Illumina; risk factor analyses were multivariable linear regression using the results of an owner-completed survey.

**Results:** 3GC-R *E. coli* were excreted by 20/303 rural and 31/297 urban dogs. Dog/human sharing was evident for the dominant canine 3GC-R sequence type, ST963(*bla*_CMY-2_). Cattle/dog sharing was evident for CTX-M-14 and CTX-M-32-producing *E. coli* from rural dogs, including sharing of plasmid pMOO-32, which is common on cattle farms in the area. Feeding raw meat was associated with carrying 3GC-R *E. coli* in rural dogs, but not in urban dogs, where swimming in rivers was a weak risk factor.

**Conclusions:** Given clear zoonotic potential for resistant canine *E. coli*, our work suggests interventions that may reduce this threat. In rural dogs, carriage of 3GC-R *E. coli*, particularly CTX-M producers, was phylogenetically associated with interaction with local cattle and epidemiologically associated with feeding raw meat. In urban dogs, sources of 3GC-R *E. coli* appear to be more varied and include environments such as rivers.

## Introduction

Rapidly rising rates of antibacterial resistance (ABR) and the slow pace of developing new antibacterial drugs make understanding the mechanisms of selection and transmission of ABR bacteria important objectives.^1,2^ *Escherichia coli* is a good example of a bacterium where framing these questions in the context of One Health research is valuable.^2,3^ There is evidence, for example, that humans are more likely to be colonised by ABR *E. coli* if they interact with certain environments or with certain animals, including domestic pet dogs.^4–6^

We have established a 50 x 50 km study area in Southwest England to investigate the transmission of ABR *E. coli* between One Health “compartments” - environments, farms, animal species and specific cohorts of animals or humans.^7–12^ Our work focusses on *E. coli* resistant to “critically important” antibacterials according to World Health Organization designations: fluoroquinolones and third-generation cephalosporins (3GCs).

Based on recent surveys of 3GC-resistant (3GC-R) *E. coli* from humans (urinary isolates)^7^ and dairy cattle (from faecal samples)^9^ within our study area, we concluded that plasmid-mediated 3GC-R is common in isolates from cattle and humans, and some plasmids are shared between these compartments.^7,9^ One 3GC-R plasmid identified in our work, named pMOO-32, was found in *E. coli* isolates representing 27 sequence types (STs) and on 25 dairy farms; it was also readily transmissible in the laboratory into *E. coli* of STs commonly found in human infections.^9^ Accordingly, it was surprising that we did not find pMOO-32 in *E. coli* from humans,^9^ but as the human isolates tested were from clinical (urine) samples,^7^ it may be that there has been insufficient time for pMOO-32 on farms to spill over into the local human population as part of the stable commensal flora, and then for this flora to cause opportunistic infections.

One of the aims of the work reported here was to investigate if pMOO-32 was present in *E. coli* within our study area in environments other than dairy farms. We chose first to investigate commensal *E. coli* from domestic pet dogs to test the hypothesis that dogs living in the vicinity of farms known to have pMOO-32-positive *E. coli* (“rural dogs”) would be more likely to carry pMOO-32-positive *E. coli* than dogs living in the city (“urban dogs”). We therefore recruited cohorts of dogs within both rural and urban portions of our study area. This led to a wider analysis of the differences between the molecular ecology of 3GC-R *E. coli* carried by urban and rural dogs as well as the behavioural risk factors associated with carriage of such *E. coli*.

## Materials and Methods

### Participant Recruitment, Ethics and Sample Collection

Ethical approval was obtained from the Faculty of Health Research Ethics Committee, University of Bristol (Ref: 89282) alongside Ethical Approval of an Investigation Involving Animals (ref: UB/19/057).

Dog owners were recruited at local dog-walking sites between September 2019 and September 2020. All gave informed consent to participate. A standardised questionnaire (**Table S1**) was provided to collect demographic data and variables chosen as being potentially associated with carriage of ABR *E. coli* in dogs. Faecal samples were provided by owners in dog-waste bags. Postal packs including a stool sample collection pot were provided for owners wanting to participate in the study but unable to provide a faecal sample at the time of recruitment. Samples were either transported to the laboratory on the day of collection or delivered through the post. Once at the laboratory, samples were refrigerated and processed within 48 h.

### *Sample Processing and Selection of 3GC-R* E. coli

A portion of each faecal sample (0.1-0.5 g) was weighed, and phosphate-buffered saline (PBS) was added at 10 mL/g before vortexing and mixing with an equal volume of 50% sterile glycerol. Twenty microlitres of each sample was spread onto Tryptone Bile X-Glucuronide agar plates (Sigma) containing cefotaxime (2 mg/L) based on EUCAST breakpoints ^13^ and incubated overnight at 37°C. The limit of detection for 3GC-R *E. coli* using this method was ~1,000 cfu/g of faeces. Putative 3GC-R *E. coli* were re-streaked to confirm resistance (taking a maximum of three isolates per sample).

### Risk Factor Analysis

Where variables had multiple choice answers ‘Never, Sometimes, Often, Very Often’ in the dog owner questionnaire (**Table S1**), ‘Never’ was collapsed to ‘No’ and all other responses were collapsed to ‘Yes’. Samples from only one dog per household were included in the analysis, with the dog chosen at random prior to data being obtained. Preliminary Chi-squared tests were used to determine associations between the binary variables and sample-level positivity (where one sample represented each dog) for 3GC-R *E. coli*. Univariable logistic regression was then performed to determine crude odds ratios between positivity for 3GC-R *E. coli* and each variable. Finally, a multivariable logistic regression model was built using a backward stepwise method and identified statistically significant (p ≤0.05) variables associated with sample-level positivity for 3GC-R *E. coli*. Hosmer-Lemeshow goodness of fit tests were used to test the fit of the final multivariable model.

### PCR and WGS

Three multiplex PCR assays were used: to identify plasmid pMOO-32,^9^ various *bla*_CTX-M_ groups, and other β-lactamase genes.^7^ At least one isolate per sample was selected for WGS with multiple isolates from the same sample being sequenced only if they produced different multiplex PCR profiles. WGS was performed by MicrobesNG as previously,^11^ and analysed using ResFinder 4.1^14^ and PlasmidFinder 2.1.^15^ The pMOO-32 plasmid reference (assession: MK169211.1) was used to map (using blastn) contigs from WGS data.

### Phylogenetics

WGS data from 3GC-R canine *E. coli* isolates were compared with data from 3GC-R human ^7,8^ or dairy farm ^8,9^ isolates. WGS data where >500 contigs were present were excluded due to relatively poor assembly. Only one isolate with the same ST and resistance gene profile for each farm, dog or human was used. Sequence alignment and phylogenetic analysis was carried out as described previously^11^ using the GTRGAMMA model of rate of heterogeneity. SNP distances were determined using SNP-dists (https://github.com/tseemann/snp-dists) and phylogenetic trees were illustrated using Microreact (https://microreact.org/).^16^ Relevant reference genomes are shown in **Table S2**.

## Results

### *Prevalence and Molecular Ecology of 3GC-R* E. coli *in Rural and Urban Dogs*

In total, 303 rural dogs from 274 households and 297 urban dogs from 289 households were recruited. The two regions were centred 32 kilometres apart. The median age of rural and urban dogs was 6.5 (range 0.5-17.5) and 5.0 (range 0.5-17) years, respectively (**Figure S1)**. 3GC-R *E. coli* excretion was detected in 6.6% (n=20) and 10.1% (n=31) of rural and urban dogs, respectively (not significantly different [χ^2^ p=0.09]), with no significant difference in the age distributions of dogs excreting or not excreting 3GC-R *E. coli* (Mann-Whitney U-Tests, p=0.11 for each cohort).

From the 20 rural dogs excreting 3GC-R *E. coli*, 21 unique isolates were selected for WGS. ST963 was the most common (4/21, 19% of isolates) of 12 STs (**Table 1**). From the urban dogs, 36 unique isolates were sequenced; ST963 again was the most common (6/36, 17% of isolates) of 18 known STs (**Table 1**).

**Table 1.** 3GC-R genes/mutations in *E. coli* isolated from rural (n=21 isolates) and urban (n=36 isolates) dog faecal samples and their STs.

|  | 3GC-R genes | Count | %<br>Isolates<br>Positive | ST (CMY-2 plasmid) |
| --- | --- | --- | --- | --- |
| **Rural dogs** | CMY-2 | 7 | 33 | 4 x ST963, 2 x ST69, ST349(p92) |
|  | CTX-M-14 | 6 | 29 | ST48, ST69<sup>†</sup>, ST88<sup>*†</sup>, ST117, ST2077<sup>‡</sup>, ST6284 |
|  | CTX-M-15 | 3 | 14 | ST10, ST38, ST48 |
|  | CTX-M-1 | 3 | 14 | 2 x ST117, ST540 |
|  | CTX-M-32 | 1 | 5 | ST10<sup>‡</sup> |
|  | *ampC*-promoter (-32T>A) | 1 | 5 | ST967 |
|  | *ampC* promoter (-42C>T) | 1 | 5 | ST88<sup>*†</sup> |
| **Urban dogs** | CMY-2 | 18 | 50 | 5 x ST963, ST963<sup>¶</sup>, 2 x ST372(p96), ST372(p92), 2 x ST2541(p96), ST2541(p92), ST973(p96), ST973(p96)<sup>†</sup>, 2 x ST1850(p92), ST46(p96)<sup>**‡</sup>, ST297(p96) |
|  | CTX-M-15 | 11 | 31 | ST58, ST58<sup>§</sup>, ST46<sup>‡</sup>, ST46<sup>**‡</sup>, ST10, ST38<sup>†</sup>, ST69, ST2852, ST3036 ST3580<sup>§</sup>, unknown ST |
|  | CTX-M-1 | 3 | 8 | ST88<sup>§</sup>, ST1584, unknown ST |
|  | CTX-M-27 | 1 | 3 | ST69 |
|  | blaSHV-12 | 1 | 3 | ST117 |
|  | *ampC* promoter (-42C>T) | 3 | 8 | ST88, ST162, ST297<sup>¶</sup> |

\*One rural isolate, ST88, harboured both CTX-M-14 and an *ampC* promoter mutation. \*\*One urban ST46 isolate had genes for CMY-2 and CTX-M-15. Symbols †, ‡, § or ¶ identify each of two rural and four urban dogs (one symbol for each), from which multiple isolates were analysed.

The predominant 3GC-R mechanism in *E. coli* from both rural and urban dogs was CMY-2 (33% and 50% of isolates, respectively; not significantly different [χ^2^ p=0.22]). In three rural and two urban ST963 dog isolates, *bla*_CMY-2_ was chromosomally inserted downstream of *nhaR*. The gene was inserted upstream of *selA* and *rhsC* in two and one urban ST963 isolates, respectively. Of the remaining CMY-2 positive isolates, eight (all urban, and covering five STs) carried *bla*_CMY-2_ on an IncI1 plasmid almost identical to p96 (Accession number CP023370.1) and five isolates (one rural plus four urban, covering four STs) carried *bla*_CMY-2_ on an IncI1 plasmid almost identical to p92 (Accession number CP023376.1). In the remaining rural CMY-2-producing isolates (both ST69), *bla*_CMY-2_ was embedded within an IncFII plasmid that had in turn inserted into the chromosome close to *ompW*.

Alongside these CMY-2 producers, chromosomal AmpC hyper-production was seen in three urban and two rural dog isolates (**Table 1**). Overall, therefore, AmpC-type mechanisms were found in 10/21 (48%) and 21/36 (58%) of 3GC-R *E. coli* excreted by rural or urban dogs, respectively, with CTX-M-type β-lactamases being responsible for 3GC-R in all remaining isolates, except one urban isolate which produced SHV-12 (**Table 1**). Of the CTX-M producers, CTX-M-14 was the most common among rural dogs (6/21, 29% of isolates) but was not seen in any urban dogs, where CTX-M-15 was most common (11/36, 31% of isolates compared with 3/21, 14% of isolates from rural dogs). CTX-M-32 was found in one rural dog and CTX-M-27 was found in one urban dog isolate (**Table 1**).

Owner survey data revealed that 70 rural and 33 urban respondents had multi-dog households, which was a statistically significant difference (χ^2^ p<0.001). Amongst the dogs for whom faecal samples were obtained, eight pairs of urban dogs and 27 pairs along with one triplet of rural dogs that shared a home were included. In 3/28 rural and 3/8 urban, sampled multi-dog households, at least one of the dogs tested positive for 3GC-R *E. coli* but in only one of these (rural) households, did two dogs test positive. WGS showed that the 3GC-R *E. coli* excreted by these two dogs were of different STs and carried different 3GC-R genes. Therefore, we found no evidence that dogs sharing a household were colonised with the same 3GC-R *E. coli*.

### *pMOO-32-like Plasmid Found in a Rural Dog* E. coli

WGS confirmed that one rural dog (Dog 118) was carrying *E. coli* ST10 encoding CTX-M-32. WGS contigs from two isolates from Dog 118 and from pMOO-32-positive cattle isolates from three dairy farms located closest to the home of Dog 118 (collected during our earlier surveillance study)^10^ were mapped (**Fig. S2**) to the archetypal pMOO-32 plasmid (accession number MK169211.1). Slight differences between the pMOO-32-like plasmid from Dog 118 and the archetypal pMOO-32 plasmid (small regions representing transposable elements absent in the dog isolates) were also seen in pMOO-32-like plasmids from the three local dairy farms, confirming that the plasmid found in Dog 118 was more similar to pMOO-32-like plasmids circulating on local farms than the pMOO-32 archetype, from a more geographically distant farm.

### *Phylogenetic Analysis of 3GC-R* E. coli *Across Four One Health Compartments*

The core genomes of 396 3GC-R *E. coli* isolates from four compartments – rural dogs, urban dogs, humans and dairy cattle – collected within our 50 x 50 km study area were aligned and a phylogenetic tree constructed (**Fig. S3**). The aim was to identify general ST overlaps and possible evidence for sharing between compartments, where pairs of isolates had <100 single nucleotide polymorphism (SNP) differences between them, which we have previously used to define sharing.^11^

ST117 isolates were found in cattle, rural dogs and one urban dog. A detailed tree revealed close relatedness between ST117 isolates from one rural dog (Dog 122) and cattle isolates from three farms with SNP distances of 42, 53 and 56, which is suggestive of sharing (**Fig. 1**). Notably, all four closely related isolates carried *bla*_CTX-M-14_ embedded in the chromosome adjacent to *nlpA*. None of the five (non-ST117) *bla*_CTX-M-14-_positive isolates from rural dogs shared this insertion site.

**Figure 1.**
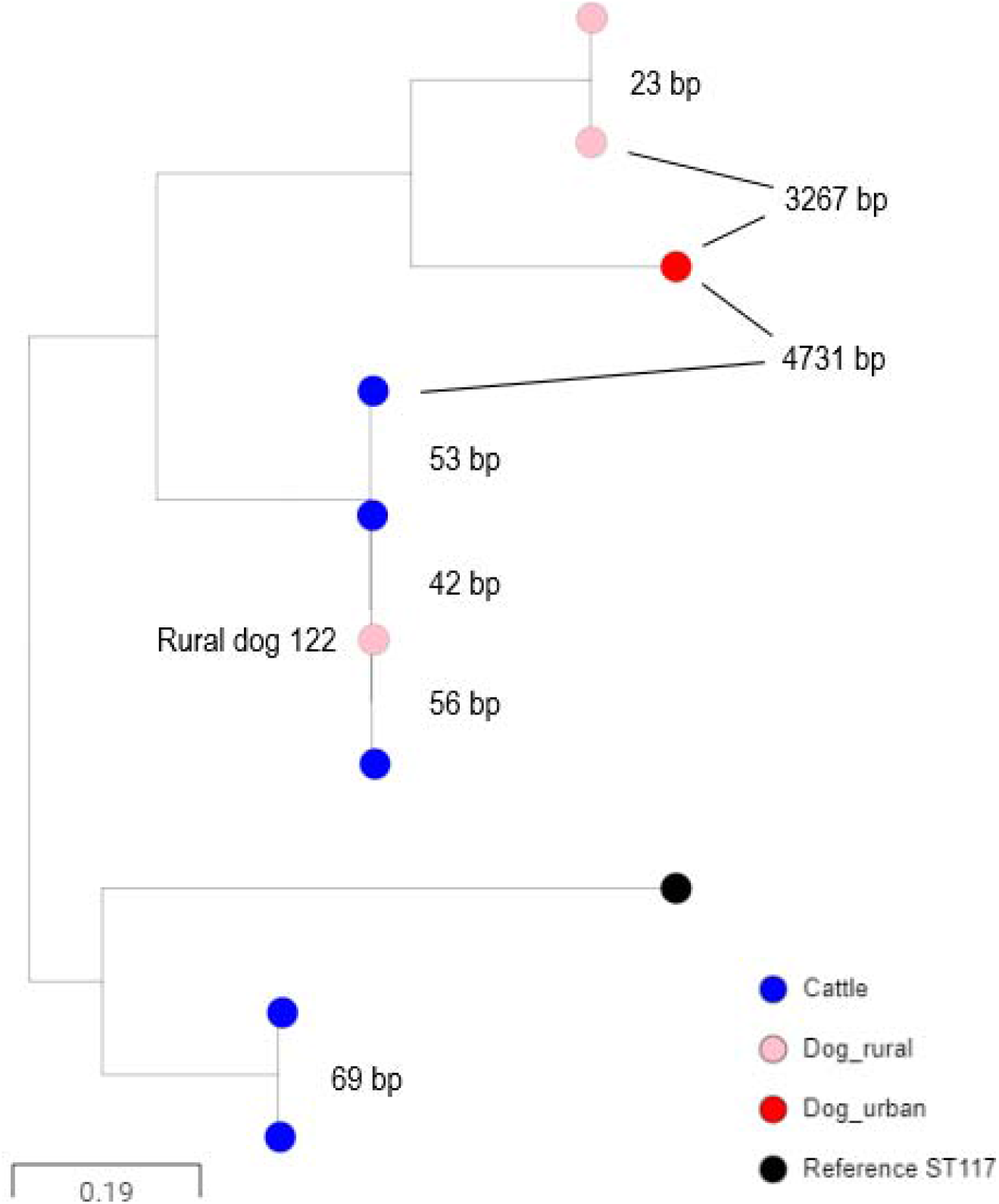
Core genome alignment of 3GC-R *E. coli* ST117 isolates of rural dogs (pink), an urban dog (red) and cattle (blue) origin. Pairwise SNP distances are labelled between isolates (bp). The isolates from Dog 122 and the three related cattle isolates carried *bla*_CTM-M-14_ embedded at the same chromosomal location.

ST58 was seen in urban dogs, humans and cattle and a detailed tree (**Fig. 2**) revealed one instance of likely dog-cattle sharing where a pair of *bla*_CTX-M-15_-positive isolates differed by 62 SNPs. In both isolates, *bla*_CTX-M-15_ was inserted into the chromosome close to *pykF*.

**Figure 2.**
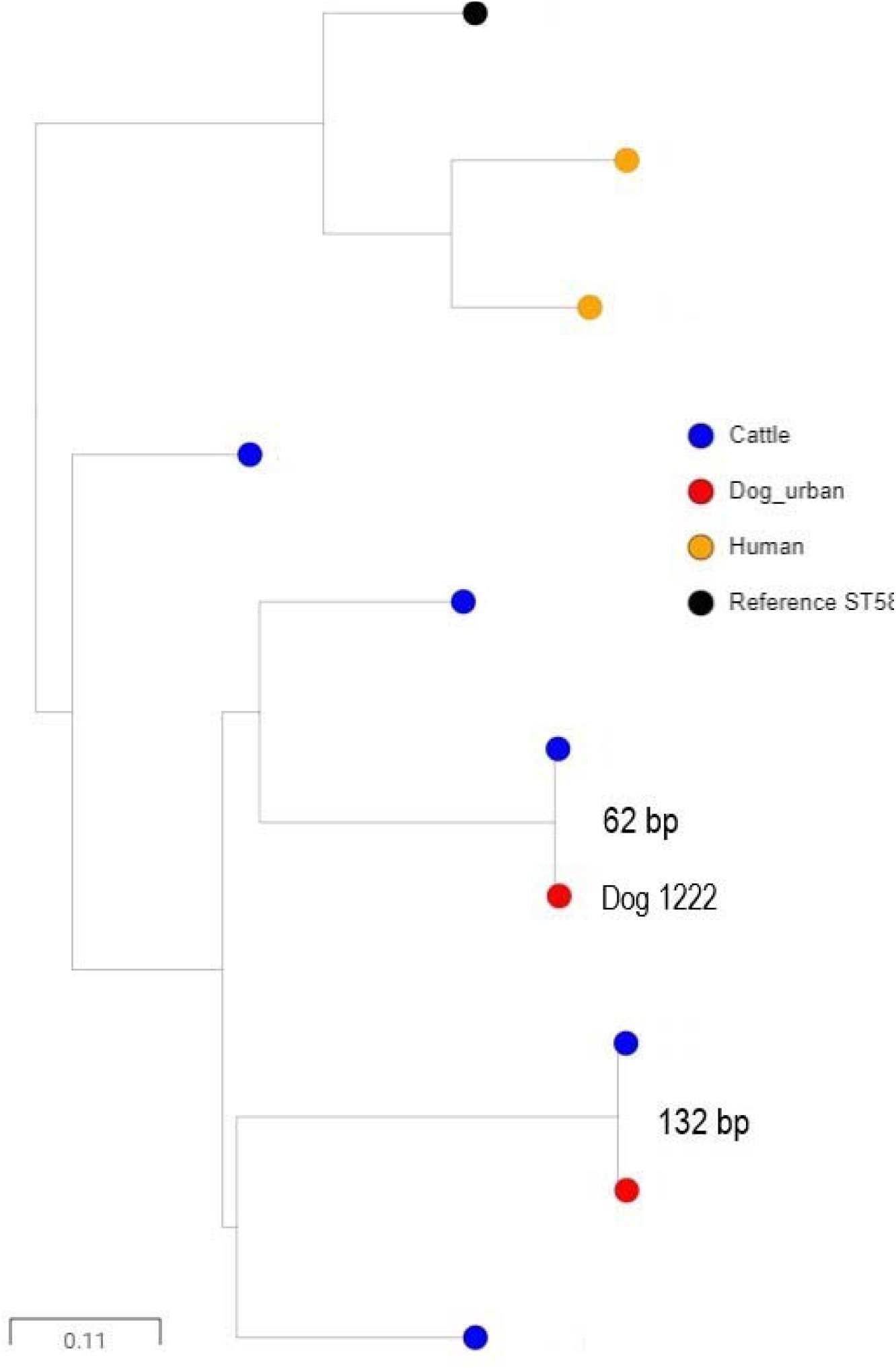
Core genome alignment of 3GC-R *E. coli* ST58 isolates of urban dog (red), cattle (blue) and human (orange) origin. Pairwise SNP distances are labelled between isolates (bp). The isolates from Dog 1222 and the related cattle isolate carried *bla*_CTM-M-15_ and *qnrS* embedded at the same chromosomal location.

ST963 isolates were seen in urban and rural dogs and humans but were not found in cattle. A detailed tree (**Fig. 3**) showed that the SNP distances between ST963 human and rural dog isolates were between 106 and 191 SNPs. Isolates from humans and urban dogs showed closer relatedness at between 64 and 107 SNPs, suggesting some sharing. Notably, the core genome phylogeny overlaid onto the *bla*_CMY-2_ genomic location. Specifically, the closely related cluster of two urban dog and two human isolates all carried *bla*_CMY-2_ inserted into the chromosome at an identical position (**Fig. 3**).

**Figure 3.**
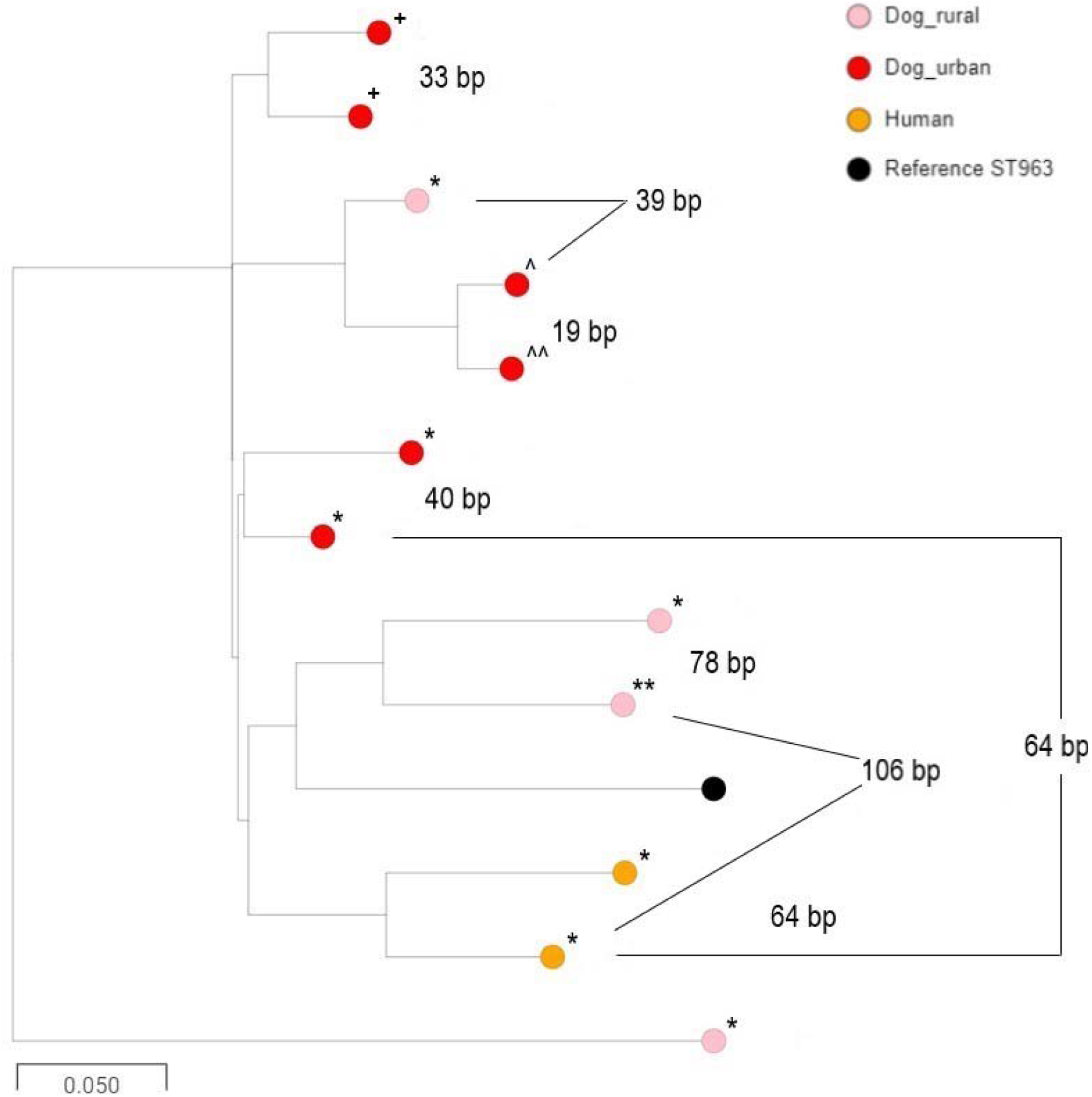
Core genome alignment of 3GC-R *E. coli* ST963 isolates of urban dog (red), rural dog (pink) and human (orange) origin. Pairwise SNP distances are labelled between isolates (bp). Chromosomal insertion points for *bla*_CMY-2_ are represented by symbols: * insertion downstream of *nhaR*; ^**+**^ inserted upstream of *selA*; **^** insertion upstream of *rhsC*. ** or **^^** indicate that we were unable to be determine location of insertion due to short read sequence assembly issues.

### Risk Factor Analysis

Preliminary χ^2^ analysis identified significant associations between excretion of 3GC-R *E. coli* and dogs being fed raw food. Additionally, and only in urban dogs, playing/swimming in rivers was found to be positively associated with excretion of 3GC-R *E. coli* (**Tables S3, S4**). In rural dogs, this association with raw feeding was observed when all raw-fed dogs were considered, and when only dogs fed commercial raw food marketed as dog food were considered (χ^2^ p<0.001 in both cases). In urban dogs, the association with raw feeding was weaker, and not significant when only feeding commercial food was considered (χ^2^ p=0.051 compared with p=0.019 when all raw-fed dogs were considered).

There was no association identified between excretion of 3GC-R *E. coli* and walking rural dogs in environments alongside cattle (**Tables S3, S4**), even when splitting the data into categories where the survey question was answered “often/very often” versus “sometimes/never” (χ^2^ *p*=0.66). There was also no association identified between rural dogs walking near cattle when we considered excretion of CTX-M or CMY-2/AmpC-producing *E. coli* separately (χ^2^ p≥0.29). Nonetheless, two out of the five rural dogs excreting CTX-M-producing *E. coli* and walked often/very often around cattle excreted *E. coli* closely related to *E. coli* from cattle in our study region: one ST117 *E. coli* producing CTX-M-14 (**Fig. 1**) and one ST10 *E. coli* producing CTX-M-32 (pMOO-32), as discussed above. In contrast, none of the six rural dogs excreting CTX-M-producing *E. coli* and walked sometimes/never around cattle excreted *E. coli* phylogenetically related to those from cattle.

Univariable followed by multivariable logistic regression concluded that raw feeding of rural dogs (but not urban dogs) was associated with increased odds of excreting 3GC-R *E. coli* (univariable Odds Ratio [OR] 9.81, 95% CI 3.56 to 27.06, p<0.001; multivariable OR 12.1, 95% CI 3.3 to 43.8, p<0.001). An increased odds of excreting 3GC-R *E. coli* in urban dogs (but not rural dogs) was also weakly associated with playing/swimming in a river (univariable OR 2.20, 95% CI 0.98 to 4.96, p=0.06; multivariable OR 2.3, 95% CI 1.0-5.2, p=0.05).

Only 26/303 (8.6%) of rural dogs and 31/297 (10.4%) of urban dogs were raw fed whereas 7/11 rural dogs and 4/12 urban dogs excreting CTX-M producing *E. coli* were raw fed. There was therefore a significant association between raw feeding and excretion of CTX-M-positive *E. coli* across both urban and rural cohorts (χ^2^ p<0.001). In contrast, only 1/9 rural dogs and 4/20 urban dogs excreting CMY-2/AmpC-producing *E. coli* were raw fed, which was not a significant association (χ^2^ p>0.15).

## Discussion

The overall aim of this work was to test the general hypothesis that, within our 50 x 50 km study area, rural dogs tend to carry 3GC-R *E. coli* that are “cattle like” and urban dogs carry those that are “human like”. Rural dogs were found to excrete CTX-M-14 and CTX-M-32-producers, which were not excreted by urban dogs and were rare or non-existent among human urinary isolates,^7^ but common in cattle.^9^ Urban dogs, in contrast, excreted CTX-M-15, CTX-M-27 and SHV-12-producers, which were rare or non-existent in rural dogs and cattle,^9^ but common in humans.^7^

Furthermore, CTX-M-32-encoding plasmid pMOO-32, which had been identified on 27 dairy farms within our study region,^9^ was found in an ST10 isolate from one rural dog. Notably, this dog – according to the owner survey – regularly walked in fields where cattle were present and was not fed raw meat. There was also good phylogenetic evidence for sharing of a CTX-M-14-producing ST117 *E. coli* circulating in the local cattle population by another dog that also regularly walked close to cattle (**Fig. 1**).

These observations provide evidence that sharing of CTX-M-producing *E. coli* between cattle and dogs may come through interaction of dogs with the near-farm environment. However, raw feeding was also shown to be associated with the excretion of CTX-M-producing *E. coli* by both rural and urban dogs. This result agrees with earlier more general observations that raw feeding is associated with excretion of resistant *E. coli* in dogs and adds to the increasing body of evidence that raw feeding poses a significant zoonotic threat.^12, 18–21^ Moreover, when considering *E. coli* core genome phylogeny, a cattle/rural dog compartmentalisation was not observed except for the two CTX-M producers discussed above. Accordingly, 3GC-R *E. coli* seen in rural dogs are likely to be predominantly from a source other than cattle reared nearby.

It is known that humans and dogs readily share *E. coli*.^4–6^ However, this study also found that *E. coli* excreted by urban dogs were not more phylogenetically related to human isolates than those excreted by rural dogs, except perhaps in the case of CMY-2-producing ST963 (further discussed below). One explanation for this finding might be that the human isolates all came from clinical urine samples.^7^ As such, factors other than ABR and transmission dynamics (including bacterial virulence and human-specific colonisation potential) may be influencing the type of isolates we cultured. It is also notable that, despite it being found in a dog, pMOO-32 has not been seen in 3GC-R urinary *E. coli* recovered from humans in our study region.^7^ However, it may take some time for commensal pMOO-32 carriage to manifest as drug-resistant opportunistic infections. CTX-M-32 is not phenotypically different from CTX-M-15,^17^ which are already dominant in human 3GC-R *E. coli* in this region,^7^ and pMOO-32 does not carry any other resistance genes clinically important for humans.^9^ So even if pMOO-32 “zoonosis” were to occur, it would be expected to have limited additional clinical impact. Observing it’s transmission, however, would clearly demonstrate how ABR plasmids can enter human populations from farmed animals – perhaps via domestic pets acting as intermediaries – and this possibility should serve to increase efforts to reduce the prevalence of such plasmids on farms.

In contrast to the above observations concerning CTX-M-producers, there was no association between raw feeding or interaction with cattle and excretion of CMY-2/AmpC-producing *E. coli* by dogs, so this suggests an alternative origin not related to near-farm environments or food. Nevertheless, CMY-2/AmpC-production collectively caused 3GC-R in approximately 50% of both rural and urban dog isolates. This approximately 50:50 (CMY-2/AmpC:CTX-M) split was also seen in a recent analysis of 3GC-R *E. coli* from dairy farms in the same study area,^9^ where amoxicillin/clavulanate use was associated with finding AmpC-mediated 3GC-R *E. coli* in farm samples.^8^ A study examining prescribing at small animal veterinary practices in the UK found that amoxicillin/clavulanate was the most common antibacterial prescribed, accounting for 36 % of prescriptions,^22^ and it has been demonstrated that routine amoxicillin/clavulanate treatment selects for increased 3GC-R *E. coli* in the faeces of dogs.^23^

Owner-reported (last 6 months) antibiotic treatment of the dogs enrolled in our study was similar in rural and urban dogs and the overall treatment incidence with antibiotics was not associated with excretion by dogs of 3GC-R *E. coli* (**Tables S3, S4**). One hypothesis, therefore, is that maternal transmission drives colonisation of dogs with CMY-2/AmpC-producing *E. coli* at a young age and this colonisation is maintained at population level by historical and ongoing selection caused by amoxicillin/clavulanate use. This may also explain why plasmids encoding CMY-2 in *E. coli* excreted by dogs in this study were almost identical to plasmids (p96 and p92) found in other UK canine cohorts.^24,25^ Importantly, CMY-2-producing ST963 (where *bla*_CMY-2_ was embedded in the chromosome at phylogeny-specific locations) was the most common individual type of 3GC-R isolate found in dogs in this study (**Table 1**). Recent sharing of 3GC-R *E. coli* ST963 between dogs and humans was also observed within our study region, with the strongest evidence of this being found in the urban environment (**Fig. 3**). Indeed, this sharing of ST963 between dogs and humans accounted for 25% of CMY-2-producing *E. coli* in the human urinary isolates in our study.^7^

It is interesting to consider why raw feeding was not (in the multivariable analysis) associated with excretion of 3GC-R *E. coli* by urban dogs, when there was a strong association in rural dogs. One explanation is that there are many variables (potential risk factors) associated with carriage of 3GC-R *E. coli* in the urban environment, and that the risk factors we explored in this study may not be independently strong enough to be identified through such observational analysis. It may be that higher population density (for humans and dogs) and greater contamination of urban environments with resistant bacteria from both are also important risk factors which we did not explore in this study. It is therefore interesting to note that river swimming was the only significant (though rather weak) risk factor associated with 3GC-R *E. coli* carriage in urban dogs and this may suggest that urban river water in our study region may be contaminated with 3GC-R *E. coli*.

In conclusion, these analyses suggest interventions that might be used to reduce 3GC-R *E. coli* carriage in dogs, and, in so doing, reduce any resultant zoonotic threat. This work also demonstrates that there are large gaps in our understanding of the complexities of AMR transmission and selection in a One Health context, particularly in the urban environment. This is important because transmission events involving humans are likely to be far more numerous in the urban environment than those happening in the better-studied farm/environment/human interface, at least in developed countries.

## Acknowledgements

We wish to thank all the dog owners who participated in this study. Genome sequencing was provided by MicrobesNG (http://www.microbesng.uk)..

## Funding

This work was funded by grant NE/N01961X/1 to M.B.A. and K.K.R. from the Antimicrobial Resistance Cross Council Initiative supported by the seven United Kingdom research councils. J.E.S. is supported by a scholarship from the Medical Research Foundation National PhD Training Programme in Antimicrobial Resistance Research (MRF-145-0004-TPG-AVISO).

## Author Contributions

Conceived the Study: J.E.S., K.K.R., M.B.A.

Collection of Data: J.E.S. supervised by M.B.A.

Cleaning and Analysis of Data: J.E.S. A.H., O.M., supervised by V.C.G., K.K.R., M.B.A.

Initial Drafting of Manuscript: J.E.S., M.B.A.

Corrected and Approved Manuscript: All authors

## Notes

### Competing Interest Statement

The authors have declared no competing interest.

