## Supplementary Data File for "Similarities and differences in molecular epidemiology of third-generation cephalosporin-resistant *Escherichia coli* carried by dogs living in urban and nearby rural settings and associated behavioural risk factors"

**^2^University of Bristol Medical School, Population Health Sciences, Canynge Hall, 39 Whatley Road, Bristol. BS8 2PS**

**^3^University of Bristol Veterinary School, Langford House, Langford, Bristol. BS40 5DU**

**Running Title: 3GC-R *E. coli* in dogs.**

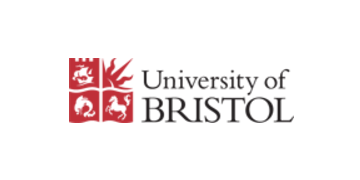
**Table S1. The Owner questionnaire**

Dog Owners Questionnaire

**Investigating the Regional Circulation of Antibiotic Resistance Genes.**

Date …………………………………

How old is your dog? …………………………………..

What breed is your dog? ............................................................................................................

1. Which types of food do you regularly feed your dog, or have they had in the past? Please tick all that apply

|  | Currently feeding | Have fed in the past |
| --- | --- | --- |
| Commercial wet food (e.g. tins or pouches) |  |  |
| Commercial dry food (e.g. kibble or pellets) |  |  |
| Commercially prepared raw diet |  |  |
| Home-prepared uncooked/raw meat |  |  |
| Home-prepared cooked meat |  |  |
| Table scraps and leftovers |  |  |
| Any other information: | | |

2. In which of these environments is your dog walked? Please tick all that apply.

|  | Never | Sometimes | Often | Very often |
| --- | --- | --- | --- | --- |
| Roads and streets |  |  |  |  |
| Parks |  |  |  |  |
| Beaches |  |  |  |  |
| Countryside, in spaces shared with livestock animals |  |  |  |  |
| Countryside but not in spaces where livestock animals are found |  |  |  |  |
| In places where cattle are kept |  |  |  |  |

**Please turn over and complete the other side**.

3. Does your dog ever swim, paddle or play in any of the following?

|  | Never | Sometimes | Often | Very often |
| --- | --- | --- | --- | --- |
| Sea or estuary |  |  |  |  |
| Lakes |  |  |  |  |
| Rivers |  |  |  |  |
| Ponds |  |  |  |  |
| Other water source (please describe briefly) | | | | |

4. Do you own any other animals? Please give brief details

5. Has your dog been unwell and therefore had antibiotics given or prescribed by your vet in the last 6 months? If so, please give brief details of the reason for antibiotic use and what was given.

6. We would be very grateful if you could give us some information about where you regularly walk your dog, particularly in the countryside. For example, you could tell us the road names, park names, map references. postcodes or nearby well-known landmarks.

**Thank you for taking the time to complete this questionnaire!**

**Table S2. Reference genomes used for sequencing alignment.**

| **Subtree** | ***E. coli* Reference Genome** |
| --- | --- |
| ST963 | Strain 746 (Assession: CP023353) |
| ST58 | Strain CFS3313 (Assession: CP026939.2) |
| ST117 | Strain MDR_56 (Assession: CP019903.1) |

**Table S3**. **Chi-square analysis of potential risk factors associated with excretion of 3GC-R *E. coli* in dogs from the rural cohort.**

Where excretion of 3GC-R *E. coli* is associated with p<0.05, p-values are presented (bold, underlined). In all other cases, p>0.05.

| **Risk factor** | **Total**  **(n=303)** | **3GC-R excreters (N=20)** |
| --- | --- | --- |
| Fed Dry food  Yes  No | 254  49 | 15  5 |
| Fed Wet food  Yes  No | 122  181 | 4  16 |
| Fed Human food  Yes  No | 164  139 | 8  12 |
| Fed Raw food  Yes  No  ***p*=** | 26  277 | 8  12  **<0.001** |
| Fed raw food in the past  Yes  No | 19  284 | 0  20 |
| Walking on streets  Yes  No | 287  16 | 19  1 |
| Walking in parks  Yes  No | 288  15 | 19  1 |
| Walking on beaches  Yes  No | 220  83 | 11  9 |
| Walking in the countryside (without livestock)  Yes  No | 256  47 | 20  0 |
| Walking in the countryside (with livestock)  Yes  No | 222  81 | 17  3 |
| Walking in countryside (with cattle)  Yes  No | 141  162 | 9  11 |
| Playing in sea estuary  Yes  No | 144  159 | 8  12 |
| Playing in lake  Yes  No | 81  222 | 5  15 |
| Playing in river  Yes  No | 165  138 | 10  10 |
| Playing in pond  Yes  No | 90  213 | 7  13 |
| Owning another dog(s)  Yes  No | 70  233 | 5  15 |
| Owning a cat(s)  Yes  No | 42  261 | 5  15 |
| Owning rodent(s)  Yes  No | 12  291 | 1  19 |
| Owning bird(s)  Yes  No | 13  290 | 1  19 |
| Owning Reptile(s)  Yes  No | 8  295 | 0  20 |
| Owning horse(s)  Yes  No | 7  296 | 0  20 |
| Owning livestock  Yes  No | 8  295 | 0  20 |
| Antibiotic use in last 6 months  Yes  No | 44  259 | 2  18 |

**Table S4. Chi-square analysis of potential risk factors associated with excretion of 3GC-R *E. coli* in dogs from the urban cohort.**

Where excretion of 3GC-R *E. coli* is associated with p≤0.05, p-values are presented (bold, underlined). In all other cases, p>0.05.

| **Risk factor** | **Total**  **(N=297)** | **3GC-R excreters (N=31)** |
| --- | --- | --- |
| Fed Dry food  Yes  No | 258  39 | 24  7 |
| Fed Wet food  Yes  No | 121  176 | 11  20 |
| Fed Human food  Yes  No | 155  142 | 17  14 |
| Fed Raw food  Yes  No  ***p*=** | 31  266 | 7  24  **0.019** |
| Fed raw food in the past  Yes  No | 21  276 | 1  30 |
| Walking on streets  Yes  No | 269  28 | 29  2 |
| Walking in parks  Yes  No | 294  3 | 31  0 |
| Walking on beaches  Yes  No | 223  74 | 25  6 |
| Walking in the countryside (without livestock)  Yes  No | 234  63 | 26  5 |
| Walking in the countryside (with livestock)  Yes  No | 190  107 | 23  8 |
| Walking in countryside (with cattle)  Yes  No | 115  182 | 9  22 |
| Playing in sea estuary  Yes  No | 174  123 | 19  12 |
| Playing in lake  Yes  No | 106  191 | 12  19 |
| Playing in river  Yes  No  ***p*=** | 162  135 | 22  9  **0.05** |
| Playing in pond  Yes  No | 136  161 | 15  16 |
| Owning another dog(s)  Yes  No | 33  264 | 2  29 |
| Owning a cat(s)  Yes  No | 51  246 | 2  29 |
| Owning rodent(s)  Yes  No | 6  291 | 1  30 |
| Owning bird(s)  Yes  No | 3  294 | 0  31 |
| Owning Reptile(s)  Yes  No | 5  292 | 1  30 |
| Antibiotic use in last six months  Yes  No | 42  255 | 5  26 |

**Figure S1. Frequency distribution of dogs in the rural and urban cohorts separated by age rounded to the nearest year.**

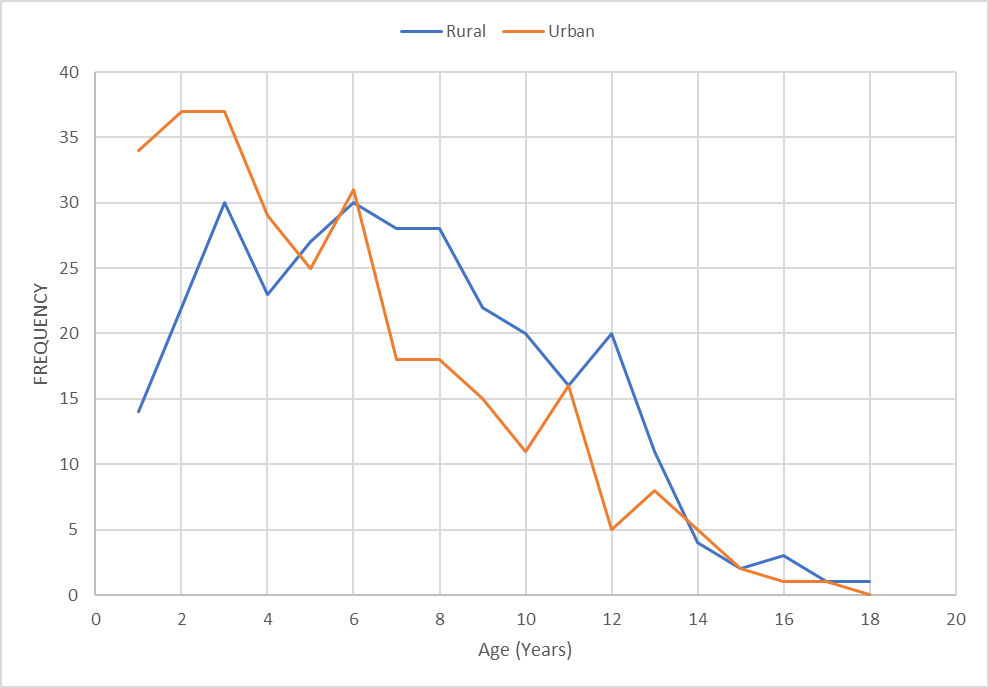

**Figure S2 Graphical representation of contig alignments for pMOO-32-like plasmids**. Contig sequences from WGS data derived from two isolates from dog 118 (dog 1 and dog 2) and isolates from three dairy farms close to the home of dog 118 (farm 1-3) were aligned with completely sequenced DUK14-2 isolate pMOO-32, from a more geographically separated dairy farm (accession number MK169211.1). Numbers indicate the same contig, with gaps in contig sequences representing insertions within the reference. To scale.

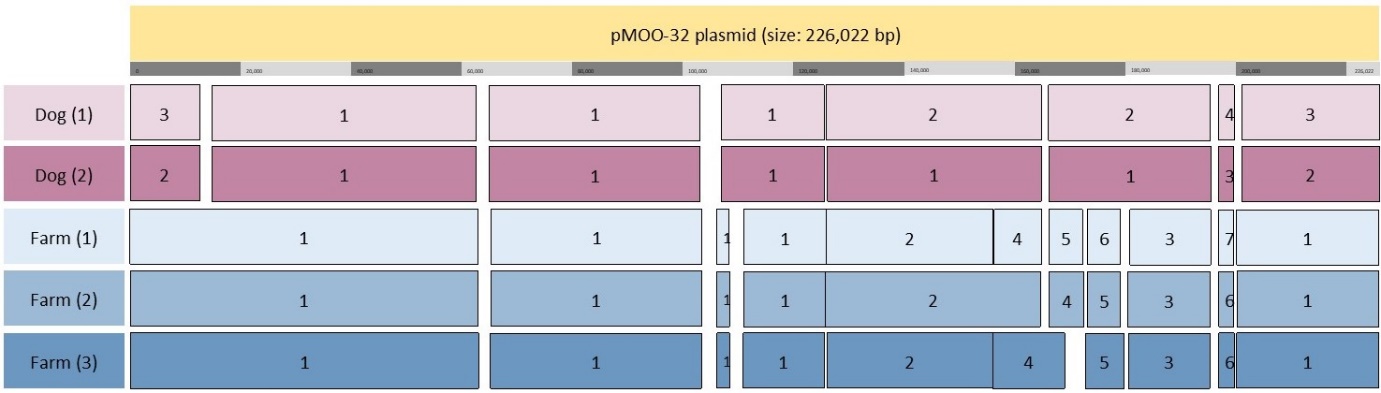

**Figure S3 Core genome phylogenetic analysis of 3GC-R E. coli from rural (pink) and urban (red) dogs, cattle (blue) and humans (orange) in our 50 x 50 km study region.** Isolates are labelled with ST.

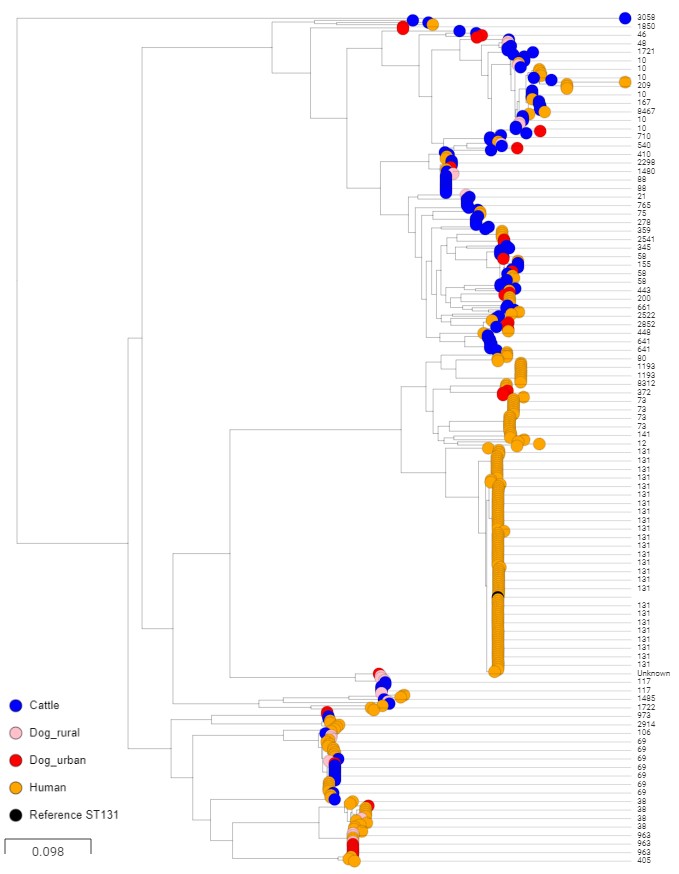
